# Tracking cortical representations of facial attractiveness using time-resolved representational similarity analysis

**DOI:** 10.1101/2020.05.21.105916

**Authors:** Daniel Kaiser, Karen Nyga

**Author notes:** Correspondence: Dr Daniel Kaiser, Department of Psychology, University of York, Heslington, York, YO10 5DD, UK.

## Abstract

When we see a face, we rapidly form an impression of its attractiveness. Here, we investigated how rapidly representations of facial attractiveness emerge in the human brain. In an EEG experiment, participants viewed 100 face photographs and rated them for their attractiveness. Using time-resolved representational similarity analysis on the EEG data, we reveal representations of facial attractiveness after 150-200ms of cortical processing. Interestingly, we show that these representations are related to individual participants’ personal attractiveness judgments, suggesting that already early perceptual representations of facial attractiveness convey idiosyncratic attractiveness preferences. Further, we show that these early representations are genuinely related to attractiveness, as they are neither explained by other high-level face attributes, such as face sex or age, nor by features extracted by an artificial deep neural network model of face processing. Together, our results demonstrate early, individually specific, and genuine representations of facial attractiveness, which may underlie fast attractiveness judgments.

## Introduction

When we see a face, we almost immediately can tell whether we find it attractive or not [1]. Beyond such first impressions, facial attractiveness affects people’s everyday lives in fundamental ways: for instance, an attractive face grants advantages in various aspects, such as increased success in dating [2,3], receiving help more often [4,5], and being more successful on the job market [6,7].

Given these varied effects of facial attractiveness, a large body of research has focused on understanding the factors that make a face attractive. One line of research has tried to establish objective physical markers of facial attractiveness [8,9], revealing that faces are more attractive when they are more similar to the average [10,11], more symmetric [11-13], or have favourable sexual characteristics [14]. This research has led into advances in computer vision, providing algorithms that can predict how attractive humans will find a particular face [15,16]. However, there is considerable agreement that there is also a subjective component to facial attractiveness, with responses varying substantially between observers [17-19]. Indeed, our impression of facial attractiveness may depend on both: an objectively attractive physical composition of visual features and an idiosyncratic appreciation of these features.

How does the brain achieve the transition from physical stimulus properties into an individual representation of facial attractiveness? To answer this question, previous studies have used event-related potentials (ERPs) obtained from EEG recordings to investigate when brain responses to more or less attractive faces differ. Although in many such studies facial attractiveness influences multiple ERP components, a key question is when facial attractiveness is *first* represented in EEG signals. The answers to this question are quite mixed. Some studies highlight relatively early ERP modulations depending on facial attractiveness, for instance at the N170 processing stage [20-26], suggesting that attractiveness is analysed during perceptual face processing. Other studies suggest that attractiveness is first analysed during the N250 processing stage [27-30], at the time when face identity is represented. Finally, some studies only find late ERP modulations, at the P3 stage or during later slow waves starting from around 350-400ms [31-37], suggesting that facial attractiveness is analysed during post-perceptual stages of cognitive processing.

Given these mixed findings, at which time do brain signals first reflect face attractiveness? One factor that complicates the interpretation of previous studies clearly is the variety in tasks (with some studies employing explicit attractiveness ratings and others employing orthogonal tasks), which makes it hard to appreciate whether ERP differences are due to variations in attractiveness or differences in task demands. Another problem is the variety in stimulus materials across studies (with different studies using faces of different genders, ages, and degrees of realism) and a lack of control for stimulus variability within individual studies, which makes it hard to assess whether ERP differences are genuinely related to differences in perceived attractiveness or other low- and high-level visual attributes of the face. Further, ERP studies can sometimes lack the sensitivity that multivariate analyses techniques offer for detecting subtle changes between conditions [38,39], thereby missing out on those temporal signatures that are weaker.

Here, we set out to resolve when cortical representations of facial attractiveness first emerge during an explicit attractiveness judgment task. By using multivariate representational similarity analysis (RSA; [40]) on EEG data recorded during this task, we were able to temporally track the emergence of representations of facial attractiveness with high sensitivity, while at the same time being able to control for other sources of variability in the faces. We found that already between 150ms and 200ms of processing, brain representations reflected facial attractiveness. We uncover three key aspects of these early representations of facial attractiveness: First, we show that even such early cortical representations are partly explained by participants’ idiosyncratic attractiveness ratings. Second, we demonstrate that cortical representations of facial attractiveness are not explained by other high-level face attributes, such as the person’s sex, ethnicity, or age. Third, we use an artificial deep neural network (DNN) to show that cortical representations of facial attractiveness are not explained by visual features used for automated face recognition. Together, our results suggest that EEG signals carry genuine and individual representations of facial attractiveness, which emerge within the first 200ms of cortical processing.

## Results

We asked participants (n=23) to rate the attractiveness of 100 male and female faces (see Figure 1a for examples). Stimuli were highly controlled full-front face photographs taken from the Face Research Lab London Set [41], eliminating many sources of visual variability. On every trial, participants saw one of the faces for 1.450ms and subsequently answered two questions (Figure 1b). On the first question, they indicated whether they found the face attractive or not (hereinafter referred to as “yes/no response”). On the second question, they indicated how attractive they found the face on a 1-7 scale (hereinafter referred to as “attractiveness rating”). We compared participants’ attractiveness ratings with ratings from a large group of observers (as provided with the Face Research Lab London Set; hereinafter referred to as “database rating”). Both the database ratings and responses collected during the experiment showed reasonable variance across the faces (Figure 1c). However, individual-participant responses were only moderately correlated with the database ratings (r=0.37 and r=0.34 for the yes/no responses and attractiveness ratings, respectively), suggesting that there was substantial inter-individual variability in attractiveness judgments.

**Figure 1.**
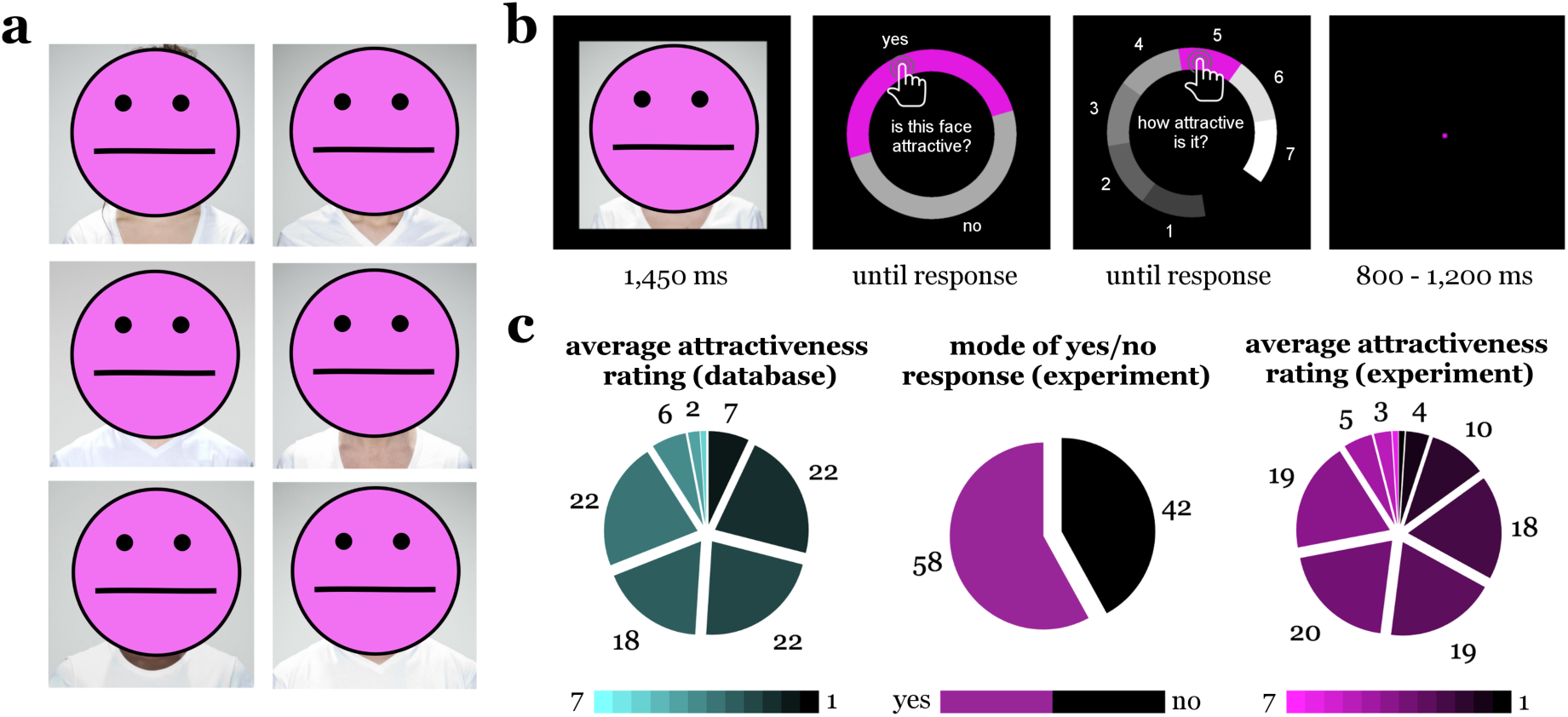
Stimuli and experimental approach. a) Stimuli were full-front and neutral face photographs of 100 individuals, covering different sexes, ages, and ethnicities. Images are taken from [41] under a CC BY 4.0 license. b) During the EEG experiment, participants viewed a single face on every trial. After seeing the face for 1,450ms, they were first asked to indicate whether they found the face attractive or not (yes/no response) and then asked to indicate how attractive they found the face on a 1-7 scale (attractiveness rating). Responses were given using the mouse. To avoid response-specific motor preparation, the positions of response options were differently arranged around a circle on every trial. c) Average attractiveness ratings from the Face Research Lab London Set database (n=2531), and yes/no responses and attractiveness ratings given by our participants (n=23). Pie charts show histograms of rating responses (left and right charts) and the number of faces rated as attractive or unattractive by the majority of participants (middle chart). These data show that there was considerable variation in ratings across faces, with a comparable amount of faces being judged as attractive or unattractive in the current experiment.

To measure how brain signals differed between faces of different perceived attractiveness, we recorded participants’ brain activity using a 64-channel EEG system. We analysed EEG signals using time-resolved multivariate representational similarity analysis (RSA; [40,42]). In RSA, the organization of brain representations is assessed by means of the pairwise (dis)similarity of each combination of stimuli, organized in neural representational dissimilarity matrices (RDMs). Neural RDMs were constructed separately for 34 discrete time bins (50ms width) across the epoch, from 250ms pre-stimulus to 1,450ms post-stimulus. For each time bin, response patterns (across all electrodes and across all time points within the 50ms time bin) evoked by each one face were correlated with response patterns evoked by each other face. RDM entries were created by subtracting these correlations from 1, so that each RDM entry reflected how dissimilar a given face was represented from another face at a given time. One such 100-by-100 RDM was constructed for each time bin (Figure 2a). Full details on the neural RDM construction can be found in the Materials and Methods section.

**Figure 2.**
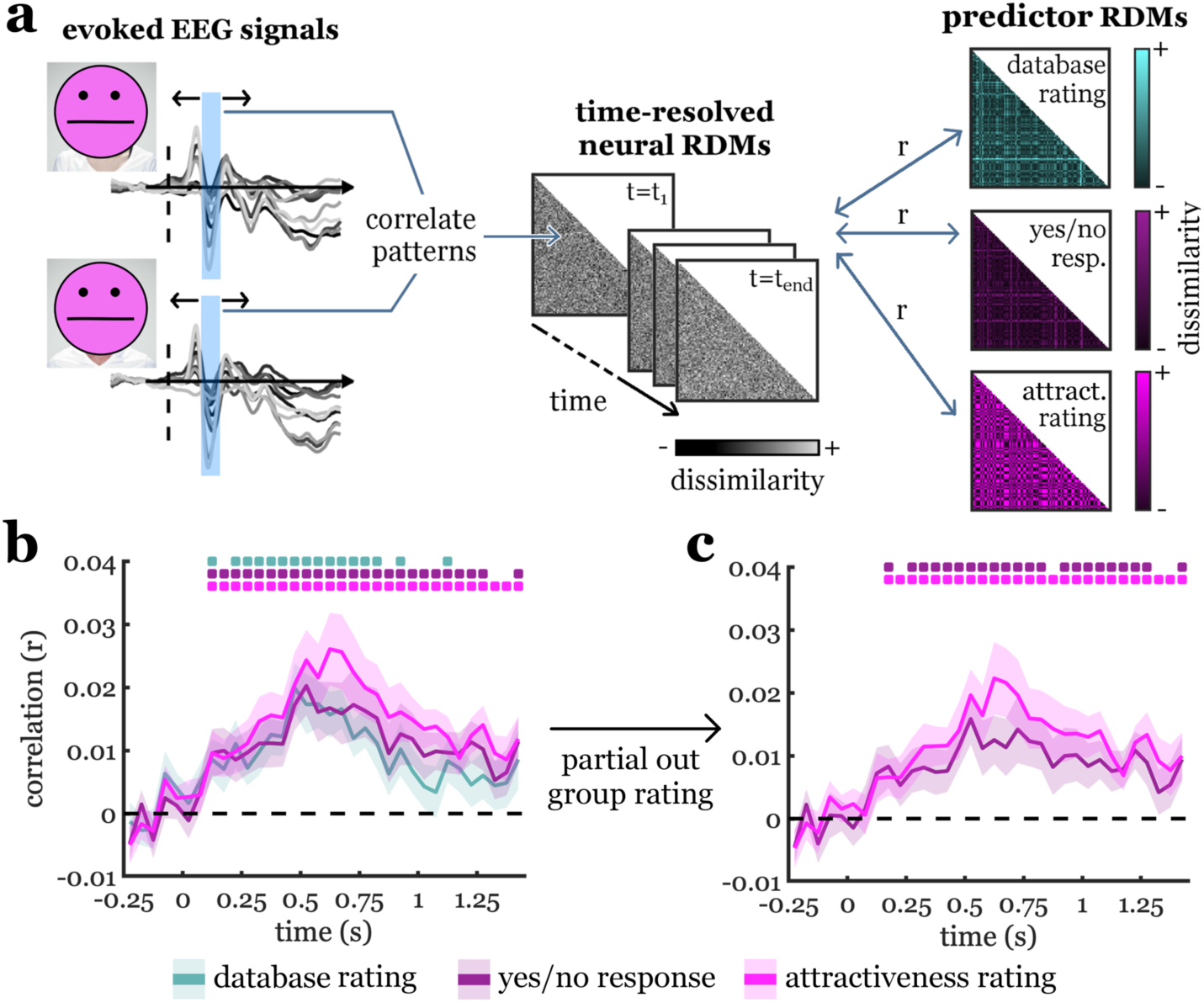
Analysis approach and key results. a) Schematic description of the representational similarity analysis. EEG response patterns for each face were extracted separately for consecutive time bins of 50ms (e.g., between 150ms and 200ms) across the epoch. Neural representational dissimilarity matrices (RDMs) were then constructed by correlating these response patterns (across time points and electrodes) for each pair of faces (for details, see Materials and Methods). This yielded 100-by-100 RDMs whose entries indexed the pairwise neural dissimilarity between faces for each time bin. For each time bin separately, neural RDMs were correlated with the three predictor RDMs, which captured the faces’ pairwise dissimilarity in attractiveness, based on (1) ratings from the Face Research Lab London Set database, (2) participants’ individual yes/no responses, and (3) participants’ individual rating responses. b) All three predictor RDMs were significantly correlated with the neural RDMs, starting from 100-150ms, suggesting an early neural representation of facial attractiveness. c) When partialing out the average database ratings, we still found that participants’ individual judgments predicted neural responses after 150-200ms, suggesting that already during this early time window, responses partly reflect an idiosyncratic signature of facial attractiveness. Error margins represent standard errors of the mean. Significance markers denote p<0.05 (corrected for multiple comparisons across time).

To investigate when cortical representations carried information about facial attractiveness, we modelled each participants’ neural RDMs using three predictor RDMs (Figure 2a): (1) a predictor RDM that reflected the faces’ pairwise dissimilarity in the average attractiveness ratings taken from the Face Research Lab London Set database, (2) a predictor RDM that reflected the faces’ pairwise dissimilarity in individual participants’ yes/no responses given during the experiment, and (3) a predictor RDM that reflected the faces’ pairwise dissimilarity in individual participants’ attractiveness ratings given during the experiment. Full details on the predictor RDM construction can be found in the Materials and Methods section.

### Early cortical representations of facial attractiveness

Correlating the predictor RDMs with the neural RDMs for each 50ms time bin across the epoch, we obtained a timeseries of how well EEG response patterns were predicted by facial attractiveness, as defined by the three predictors. Information about facial attractiveness was obtained for widespread temporal clusters across the EEG epoch and for all three types of predictors (Figure 2b): the database ratings predicted neural representations from the 100-150ms time bin (peaking at 450-500ms, peak t[22]=5.08, p<0.001, p_corr_<0.001), individual yes/no responses predicted neural representations from the 100-150ms time bin (peaking at 500-550ms, peak t[22]=4.37, p<0.001, p_corr_=0.002), and individual attractiveness ratings also predicted neural representations from the 100-150ms time bin (peaking at 600-650ms, peak t[22]=4.53, p<0.001, p_corr_<0.001). As expected, the more fine-grained attractiveness ratings predicted variance in the neural data beyond the yes/no responses (see Supplementary Figure S1a). Together, these results suggests a neural signature of facial attractiveness that already emerges during perceptual face processing.

### Early representations of facial attractiveness reflect individual attractiveness judgments

Are neural representations of facial attractiveness explicable by average preferences emerging across a large group of observers, potentially driven by a fixed set of physical face properties, with inter-individual variability reflecting merely noise? Or do they partly reflect idiosyncratic preferences, that is an individual person’s unique aesthetic preference for particular faces? To resolve this question, we performed an analysis where we modelled neural RDMs as a function of individual yes/no responses and attractiveness ratings, while controlling for the database ratings using partial correlation analysis [43-45] (Figure 2c). We found that individual attractiveness judgments still significantly predicted cortical representations, both when considering yes/no responses (from the 150-200ms time bin; peaking at 500-550ms, peak t[22]=3.90, p<0.001, p_corr_=0.012) and when considering attractiveness ratings (from the 150-200ms time bin; peaking at 600-650ms, peak t[22]=3.87, p<0.001, p_corr_=0.002). Similar results were obtained when partialing out the average judgments obtained across participants in the current experiment (see Supplementary Figure S2b). Finding that already in the 150ms-200ms time bin participants’ individual judgments explained neural responses better than group-average ratings alone suggest that even early representations of facial attractiveness are linked to individual face preferences, rather than to physical features that similarly determine attractiveness for all observers. Further, to solidify that these early representations of facial attractiveness are indeed somewhat independent from particular physical face attributes, we performed two control analyses.

### Early representations of facial attractiveness are not explained by other high-level face attributes

In the first analysis, we tested whether other high-level face attributes (the depicted person’s sex, ethnicity, or age) could explain the different cortical responses to faces of different attractiveness. To do so, we modelled neural RDMs using three predictor RDMs (Figure 3a): (1) a predictor RDM that modelled the faces dissimilarity in sex, (2) a predictor RDM that modelled the faces dissimilarity in ethnicity, and (3) a predictor RDM that modelled the faces dissimilarity in age; for details, see Materials and Methods. Each of these high-level attributes predicted parts of the neural representation across time (Figure 3b). Sex predicted face representations from the 100-150ms time bin (peaking at 350-400ms, peak t[22]=5.31, p<0.001, p_corr_<0.001), ethnicity predicted representations from the 100-150ms time bin (peaking at 100-150ms, peak t[22]=6.82, p<0.001, p_corr_<0.001), and age predicted representations from the 350-400ms time bin (peaking at 500-550ms, peak t[22]=3.68, p<0.001, p_corr_=0.007). We then modelled neural representations as a function of individuals yes/no responses and attractiveness ratings, while controlling for all three high-level attributes using partial correlation analyses (Figure 3c). We found that representations of facial attractiveness were not explained by the depicted persons’ sex, ethnicity, and age: Individual yes/no responses still predicted cortical representations from the 150-200ms time bin (peaking at 500-550ms, peak t[22]=3.70, p<0.001, p_corr_=0.009) and individual attractiveness ratings predicted representations from the 150-200ms time bin (peaking at 600-650ms, peak t[22]=4.37, p<0.001, p_corr_<0.001).

**Figure 3.**
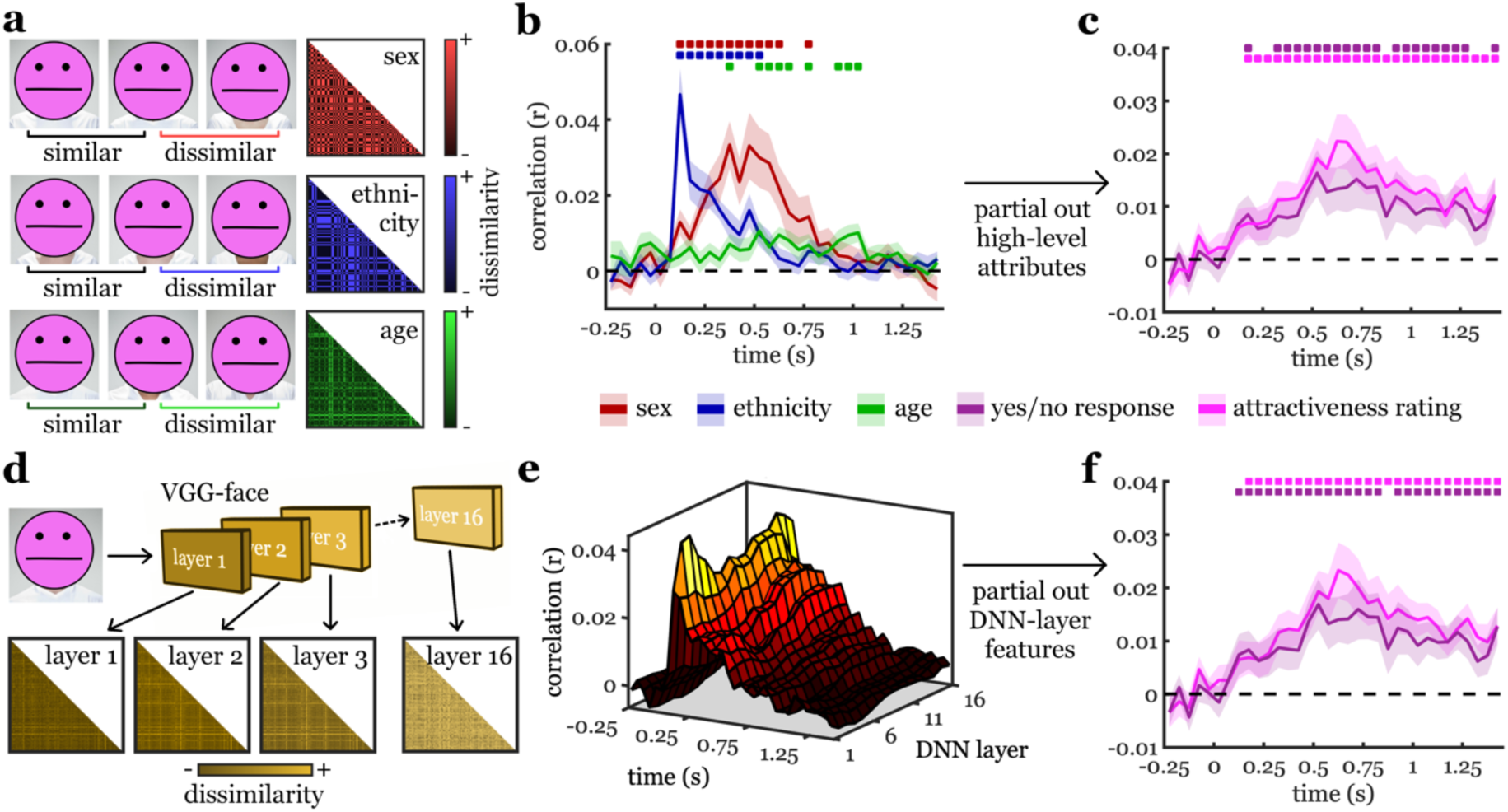
Controlling for high-level face attributes and DNN features. a) To quantify high-level face attributes, we constructed RDMs based on the faces’ dissimilarity in sex (same sex: similar, different sex: dissimilar), ethnicity (same ethnicity: similar, different ethnicity: dissimilar), and age (dissimilarity: absolute age difference between face pairs). b) The three high-level attributes all explained significant proportions of the neural representation. c) When controlling for the three attributes, participants’ attractiveness judgments still predicted neural representations from 150-200ms after onset. d) To quantify DNN features, we extracted RDMs based on the dissimilarity of DNN activation patterns for each layer of a 16-layer DNN trained on face recognition (VGG-face; see Materials and Methods). e) DNN features at each layer predicted a significant proportion of the neural representation, with early layers better predicting early activations, and late layers better predicting later activations. For detailed results, see Supplementary Figure S2. f) When controlling for both the DNN features and the high-level face attributes, we again found that after 150-200ms participants’ attractiveness judgments still predicted neural representations. Error margins represent standard errors of the mean. Significance markers denote p<0.05 (corrected for multiple comparisons across time).

### Early representations of facial attractiveness are not explained by deep neural network features

In the second control analysis, we controlled for a variety of visual features analysed during face recognition by a deep neural network (DNN). DNNs are the current state-of-the-art for modelling visual representations emerging at different processing stages of biological vision [46-48]. Here, we used a DNN trained on recognizing faces (VGG-face; [49]), which has previously been shown to accurately approximate features analysed during cortical face processing [50]. We extracted predictor RDMs from the 16 convolutional layers of the DNN, in which each entry reflected the pairwise dissimilarity (1-correlation) between the layer-specific activation vectors for two faces (Figure 3d). These RDMs predicted cortical activations starting from the 50-100ms time bin and, in line with previous reports [51-53], early representations (e.g., at 100-150ms post-stimulus) were better captured by early DNN layers, while late representations (e.g., at 250-300ms post-stimulus) were better captured by late DNN layers (Figure 3e)^1^. Next, we again modelled neural representations as a function of individual yes/no responses and attractiveness ratings, but now controlling for features extracted in each of the 16 DNN layers (and additionally for the faces’ sex, ethnicity, and age) using partial correlation analyses (Figure 3f). Critically, we again found that both yes/no responses (from the 100-150ms time bin; peaking at 500-550ms, peak t[22]=3.54, p<0.001, p_corr_=0.007) and attractiveness ratings (from the 150-200ms time bin; peaking at 600-650ms, peak t[22]=4.45, p<0.001, p_corr_<0.001) still predicted neural representations. Together, the two control analyses underscore the notion that early representations of facial attractiveness are independent from specific and fixed physical face properties.

In sum, our study reveals genuine and individually specific representations of face attractiveness. Across all analyses, these representations first emerged between 150ms and 200ms post-stimulus. This suggests that already during perceptual stages of face analysis, the brain computes how attractive we find a particular face.

## Discussion

The current study used time-resolved representational similarity analysis on EEG data to track the dynamic emergence of cortical representations related to facial attractiveness. As the key result, we demonstrate that - across multiple analyses - robust neural representations of facial attractiveness emerge after 150-200ms of vision. This timing suggests that differences in face attractiveness are related to processing differences in the N170 range [54-56]. Indeed, when comparing ERPs evoked by faces that were either rated as attractive or unattractive, we also found differences in N170 peak amplitudes, with stronger amplitudes for less attractive faces (see Supplementary Figure S3). Our results thereby support earlier studies that have shown N170 modulations as a function of facial attractiveness [20-26]. Further, this timing of attractiveness-related responses is consistent with functional neuroimaging studies showing that facial attractiveness is represented in regions of the visual face processing network [57-61]. This suggests that face attractiveness is computed during perceptual processing, at the same time when basic facial configurations are analysed [55,56,62]. Our results thereby suggest that facial attractiveness is derived from perceptual face features, potentially related to favourable face configurations.

However, given that we found representations of facial attractiveness during perceptual processing, one could argue that they are not genuine representations of attractiveness, but representations of visual features that co-vary with attractiveness in our stimulus set. Our results offer two principal refutations of this argument. First, we show that early representations of facial attractiveness are neither explained by other high-level face attributes nor by features extracted by a DNN trained on face recognition. This shows that there is no straightforward mapping between orthogonal visual features and the features used to determine facial attractiveness. Second, we show that early representations of facial attractiveness are partly explained by participant’s individual attractiveness judgments. This indicates that even such early processing of facial attractiveness is not strictly determined by the presence of particular physical face attributes: If representations of facial attractiveness were indeed a consequence of the presence or absence of a fixed set of visual features, one would expect that they are best predicted by the more stable average attractiveness rating across many observers. However, we find that representations of facial attractiveness are partly predicted by participants’ individual attractiveness judgments, suggesting that they are not directly explained by the same physical properties for all observers.

The early representation of such personal attractiveness preferences for faces suggests that differences in aesthetic appreciation are related to perceptual computations differing between individuals. Indeed, while differences in later brain representations can be attributed to task-related differences in cognitive and attentional processes, early perceptual representations are less sensitive to such processes [63,64]. Our findings therefore suggest that at the N170 processing stage individual features are weighted in idiosyncratic ways to give rise to an individual representation of facial attractiveness. This idea is in line with recent work showing that representations in object-selective visual cortex are best modelled by each person’s idiosyncratic judgments about the objects’ semantic similarities [65,66], showcasing that even perceptual brain representations can have an idiosyncratic component. Clearly, additional work needs to be done to map out this idiosyncratic component in facial attractiveness representations.

Beyond revealing an early representation of facial attractiveness, we show that EEG responses carried a sustained neural attractiveness signal. These sustained effects are in line with the later modulations of facial attractiveness stressed by some EEG studies^2^ [27-37] and with fMRI activation differences in frontal regions as a function of facial attractiveness [57-61,67]. At such later processing times, brain signals may reflect the cognitive processing of aesthetic quality: Some studies have suggested that such processes span frontal areas associated with stimulus valuation, such as the ventromedial prefrontal cortex and orbitofrontal cortex [61,67], while others have localized them to the default-mode network, potentially due to the self-referential character of aesthetic appreciation [68,69]. What is interesting is that such late processes seem to reflect aesthetic quality more generally [61,67,69], which can in principle arise from different visual inputs, such as human faces, visual scenes, and abstract stimuli, or even from non-visual inputs. At this point, to understand precisely when and where such general representations of aesthetic quality emerge, additional studies need to combine diverse stimulus sets with spatially and temporally resolved neural recordings.

In sum, our study provides evidence for an early representation of facial attractiveness, that is both genuine (i.e., unrelated to other visual face attributes) and individually specific (i.e., partly explained by participants’ personal attractiveness preferences). Finding that this representation emerged within 200ms of vision provides a neural basis for rapid judgments of facial attractiveness in real-life contexts.

## Materials and Methods

### Participants

Twenty-four adults (mean age 19.8, SD*=*1.6; 20 female) participated in the study^3^. All participants had normal or corrected-to-normal vision. Participants were psychology students at the University of York and received course credits for participation. Prior to the experiment, participants provided written informed consent. All procedures were approved by the ethical committee of the Department of Psychology at the University of York and were in accordance with the Declaration of Helsinki. One participant was excluded because of technical problems that caused missing data, leaving 23 complete datasets for analysis.

### Stimuli

The stimulus set consisted of 100 neutral full-front face photographs (for examples see Figure 1a), taken from the Face Research Lab London Set [41]. These stimuli were standardized colour photographs, limiting the variation in low-level features that are unrelated to differences in individual faces. For each face, the stimulus set includes attractiveness ratings from a large set of observers (n=2531), with considerable variance in attractiveness ratings between faces (see Fig. 1c). Each of the faces also comes with additional metadata, including the photographed person’s self-reported gender, age, and ethnicity.

### Experimental paradigm

During the EEG experiment, participants completed a single session of 700 trials, which was split into 7 blocks of 100 trials. On each trial, participants viewed one of the faces (7°-by-7° visual angle) for 1,450ms (Fig. 1b). Within each block, each face was shown exactly once, with trial order fully randomized within the block. After seeing the face, a blank screen was shown for 100ms before participants were asked two questions. On the first question, they were asked to indicate whether they found the face attractive or not by selecting yes or no. On the second question, they were asked how attractive they found the face by selecting a value between 1 (very unattractive) and 7 (very attractive). Answers to these questions were given using the computer mouse; participants could correct their answers as often as they wanted before proceeding by pressing the spacebar. To avoid response-specific motor preparation, the different response alternatives were placed at different positions around a circular response screen, as shown in Fig. 1b. Trials were separated by an inter-trial interval randomly varying between 800ms and 1,200ms. Participants were instructed to keep central fixation during the inter-trial interval and the stimulus presentation; during these periods a pink fixation dot was overlaid in the centre of the screen. Further, participants were asked to restrict eye blinks to the period when they selected their responses. Stimuli were presented on a VIEWPixx display with a 1920-by-1020 resolution and stimulus presentation was controlled using the Psychtoolbox [70].

### EEG recording and preprocessing

EEG signals were recorded using an ANT Waveguard 64-electrode system. Electrodes were arranged in accordance with the standard 10-10 system. EEG data were recorded at 250Hz sampling rate using the ANT Neuroscan Software. Offline preprocessing was performed using FieldTrip [71]. EEG data were referenced to the Fz electrode (which was discarded after preprocessing), epoched from -500ms to 1,900ms relative to stimulus onset, and baseline-corrected by subtracting the mean pre-stimulus signal for each electrode. A band-pass filter was applied to filter out 50Hz line noise. Channels and trials containing excessive noise were removed based on visual inspection. On average, 5.1 channels (SD=2.2) and 73.4 trials (SD=30.3) were removed. Blinks and eye movement artifacts were removed using independent component analysis and visual inspection of the resulting components. After preprocessing, EEG epochs were cropped from -250ms pre-stimulus to 1,450ms post-stimulus.

### Measuring neural representational similarity

To track face representations across time, we used representational similarity analysis (RSA; [40]). First, we created time-resolved neural representational dissimilarity matrices (RDMs), which reflected the pairwise dissimilarity of the faces’ brain representations across processing time. Second, we compared the organization of the neural RDMs to a set of predictor RDMs, which captured different dimensions on which the faces’ were similar or dissimilar.

Neural RDMs were constructed separately for each participant, using the CoSMoMVPA toolbox [72]. RDMs were created for 34 time bins of 50ms width each, ranging from -250ms to 1450ms relative to stimulus onset. All following analyses were done separately for each of these 34 time bins. For each time bin, we extracted a response pattern across 12 time points (covering 50ms at 250Hz) and 63 electrodes^4^. These data were then unfolded into a 756-element vector for further analyses. Before RDM construction, we performed principal-component analyses (PCAs) to reduce the dimensionality of these response vectors [39,73]. We split the available data into two independent subsets, with an equal number of trials per condition randomly assigned to each subset. The first subset of the data was used to perform the PCA decomposition. The PCA decomposition was then projected onto the second subset, retaining only the components needed to explain 99% of the variance in the first subset (99.1 components on average, SD across time: 13.2, SD across participants: 32.o). RDMs were constructed from the second subset. We first averaged across all available trials for each condition, and then correlated the response vectors for each pairwise combination of faces. These correlations were subtracted from 1 and entered into a 100-by-100 RDM. Each off-diagonal entry in this RDM thus reflected a measure of neural dissimilarity for a specific pair of faces; RDM diagonals were always empty. This procedure was then repeated with the two subsets swapped. Finally, the whole aforementioned analysis was repeated 50 times, with trials assigned randomly to the two subsets each time; RDMs were averaged across all repetitions, yielding a single RDM for each time bin.

### Tracking neural representations of facial attractiveness

To characterize the representational organization obtained from the neural signals, we compared the neural RDMs to a set of predictor RDMs, separately for each time point and participant. Like the neural RDMs, each predictor RDM contained 100-by-100 entries, which reflected the dissimilarity of pairs of faces on a particular dimension.

To assess how strongly neural representations are determined by facial attractiveness, we created three attractiveness predictor RDMs: (1) An RDM based on individual participants’ yes/no responses on the first question (“is this face attractive?”). For this RDM, pairwise matrix entries consisted of the absolute difference between yes/no-responses (coded as 1 and 2) given to the two faces. Note that participants could give different yes/no responses across repetitions of the same face. (2) An RDM based on individual participants’ attractiveness ratings on question two (“how attractive is it?”). For this RDM, pairwise matrix entries consisted of the absolute difference between attractiveness ratings (on a 1-7 scale) given to the two faces. (3) An RDM based on the average attractiveness ratings (n=2531) taken from the Face Research Lab London Set (see Stimuli). For this RDM, pairwise matrix entries also consisted of the absolute difference between attractiveness ratings (on a 1-7 scale) given to the two faces. Note that RDMs (1) and (2) were different for each participant, as they were based on their individual yes/no responses and attractiveness ratings, respectively, whereas RDM (3) was the same for every participant, reflecting a “ground truth” estimate of attractiveness ratings shared across a large group of people.

RDMs constructed from individual participants’ yes/no responses were substantially correlated to RDMs constructed from participants’ attractiveness ratings: correlations computed for each individual participant were r=0.68 on average, and the correlation between the RDMs averaged across participants was r=0.85. RDMs constructed from participants’ individual responses in the experiment only showed a weaker correlation to the RDM constructed from the average group rating in the database (average r=0.37 and r=0.34 for the yes/no responses and attractiveness ratings, respectively). When averaging the RDMs across participants, we found a much stronger correlation between RDMs reflecting the average individual-participant responses and the RDM reflecting the average ratings from the database (r=0.80 and r=0.73 for the yes/no responses and attractiveness ratings, respectively), suggesting that (1) there is a substantial amount of inter-individual variability in attractiveness ratings, and (2) average ratings in our experiment converged towards the average database ratings.

To quantify the correspondence between each of the predictor RDMs and the neural data, we computed Spearman-correlations between the neural RDMs and the predictor RDMs, separately for each participant; for these correlations only the lower off-diagonal elements of each RDM were used and the diagonal was always discarded. To establish correspondences between the neural RDM and a specific predictor RDM while controlling for other RDMs (see below), we used partial correlations [43-45]. All correlations were Fisher-transformed before entering them into statistical analyses.

### Controlling for high-level face attributes

To test whether representations were uniquely attributable to face attractiveness, we explicitly controlled for a set of high-level face properties. Specifically, we controlled for a set of person attributes that may influence attractiveness ratings: a person’s sex, ethnicity, and age. For each of these attributes we used the self-report metadata included in the Face Research Lab London Set (see Stimuli). From these data, we constructed three RDMs: (1) An RDM based on the depicted person’s sex. For this RDM, pairwise matrix entries were marked as similar when both faces were of the same sex and as dissimilar when both faces were of different sexes. (2) An RDM based on the depicted person’s ethnicity; as most of our participants (21/23) were Caucasian, ethnicity was binarized into Caucasian and non-Caucasian. For this RDM, pairwise matrix entries were marked as similar when both faces were of the same ethnicity and as dissimilar when both faces were of different ethnicities. (3) An RDM based on the depicted person’s age. For this RDM, pairwise matrix entries reflected the absolute age difference between the two faces.

### Controlling for deep neural network features

To test whether neural representations related to face attractiveness were explicable by visual features typically extracted during face processing, we used a deep neural network (DNN) model trained on face recognition. DNNs are currently the state-of-the-art models for approximating the visual feature organization emerging during perception and have been shown to accurately approximate the organization of both low-level and high-level features in visual cortex [46-48,51-53]. Here, we used a 16-layer DNN for face recognition that was pre-trained on a huge face dataset (VGG-face; [49]), as implemented in MatConvNet [74]. This DNN has previously been shown to mimic the organization of visual feature representations during cortical face processing [50]. To quantify the visual feature organization emerging in the DNN, we first ran the 100 face images through the DNN and obtained activation vectors for each of the 16 DNN convolutional DNN layers. We then constructed a model RDM for each layer by computing the representational distance (1-correlation) among each possible pair of faces.

### Statistical testing

We used one-sided t-tests against zero to identify significant correlations between the neural RDMs and predictor RDMs. False-discovery-rate (FDR) corrections were used to control for multiple comparisons across time.

## Data Availability

Stimuli and data are publicly available on OSF [75]. Other materials are available upon request.

## Acknowledgements

D.K. is supported by a Deutsche Forschungsgemeinschaft (DFG) grant (KA4683/2-1).

## Author Contributions

D.K. designed the study. D.K. and K.N. acquired data. D.K. and K.N. analyzed data. D.K. interpreted results. D.K. prepared figures. D.K. drafted the manuscript. D.K. and K.N. edited and revised the manuscript.

## Footnotes

1 Detailed results for all DNN layers can be found in Supplementary Figure S2.

2 These studies may not have found earlier differences related to facial attractiveness for varied reasons: Apart from the lower sensitivity offered by univariate ERP analyses, individual studies used largely orthogonal tasks, had small sample sizes, or did not specifically look for N170 differences.

3 This sample size was comparable with the sample sizes of previous EEG studies on facial attractiveness [20-37], who tested a median of 20 participants, and an average of 24.6 participants.

4 As during preprocessing electrodes were removed, electrode counts were lower for individual participants.

