## Supplementary Information for "Tracking cortical representations of facial attractiveness using time-resolved representational similarity analysis"

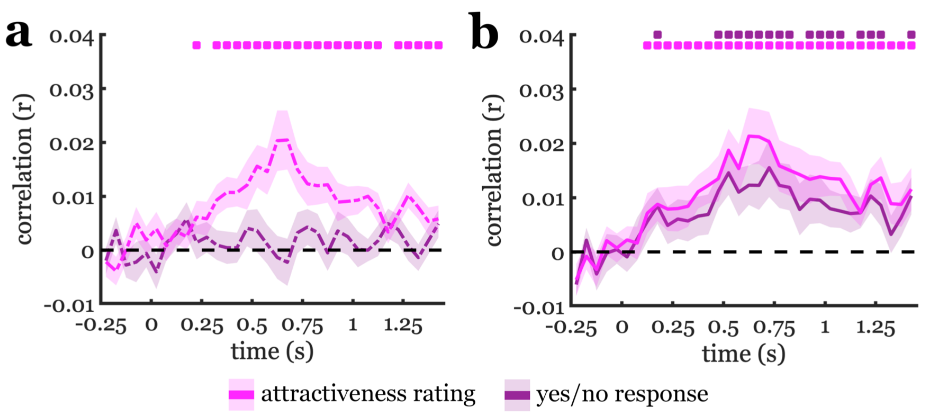


*Figure S1. Further analysis of information shared between attractiveness judgments and neural representations.* a) Comparing the information shared between neural representations and the yes/no responses and attractiveness ratings. When partialing out individual participants' yes/no responses, their individual attractiveness ratings still significantly predicted neural representations, suggesting that the attractiveness ratings add fine-grained information beyond the yes/no responses. By contrast, partialing out the attractiveness ratings did remove the correlation between yes/no responses and the neural data. b) Controlling for average attractiveness judgments in the experiment. When partialing out the average yes/no responses and attractiveness ratings (across all participants in our experiment) from participants' individual judgements, we found that individual judgments still predicted cortical representations, from 150-200ms and from 100-150ms for the yes/no responses and attractiveness ratings, respectively. This result supports the conclusion that early representations of facial attractiveness to some degree are individually specific. Error margins represent standard errors of the mean. Significance markers denote $p<0.05$ (corrected for multiple comparisons across time).

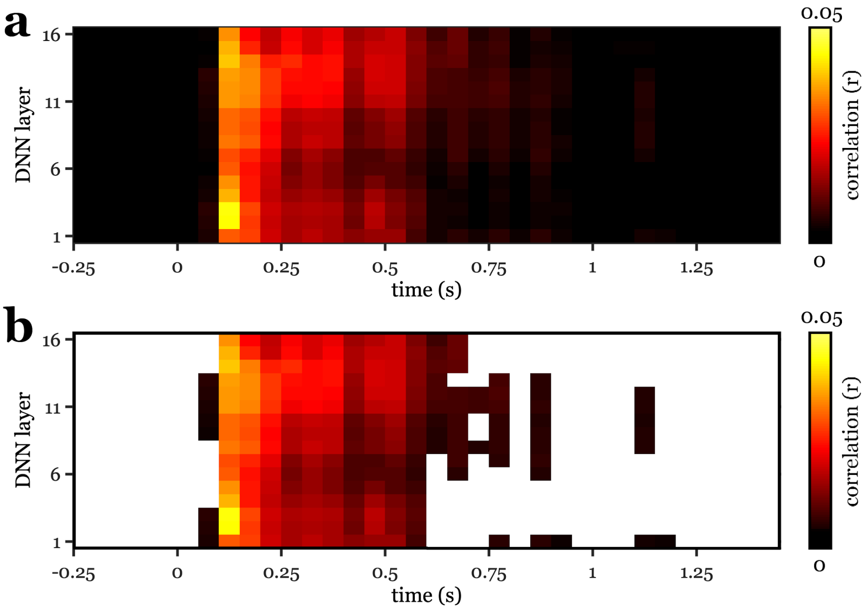


*Figure S2. Correspondence between DNN features and brain representations.* a) Correlation of RDMs extracted from each DNN layer and neural RDMs at each time point. b) Same as (a), with correlations thresholded at $p<0.05$ (corrected for multiple comparisons across time). Notably, early layers show relatively better correspondence with early brain representations (e.g., at 100-150ms after onset), while later DNN layers show relatively better correspondence with later brain representations (e.g., at 250-300ms after onset).

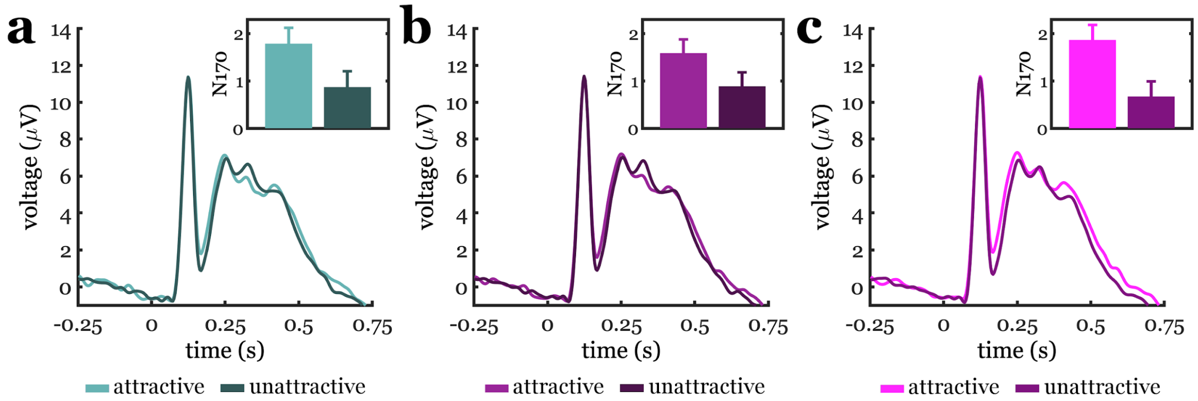


*Figure S3. N170 event-related potential results.* ERPs were averaged for electrodes P8/PO8 and P7/PO7. We then compared these ERPs to attractive faces and unattractive faces. This was done in three ways: (a) Comparing responses to faces whose database rating was above or below the median database rating, (b) comparing responses to faces where individual participants responded with yes or no on the attractiveness response (if responses were inconsistent across repetitions, we considered the more frequently chosen response), and (c) comparing responses to faces whose individual attractiveness rating was above or below the median of the respective participant's ratings. Inlays show peak N170 voltages (at 168ms post-stimulus) for the attractive and unattractive faces. For all three conditions, we found a significant N170 voltage difference, with a stronger N170 amplitude (i.e., lower voltage) for the less attractive faces (database ratings: $t[22]=2.77$, $p=0.011$; yes/no responses: $t[22]=2.39$, $p=0.026$; attractiveness ratings: $t[22]=3.74$, $p=0.001$). Error bars represent standard errors of the difference. Note that for the ERP analyses an additional low-pass filter at 30Hz was applied to the data.
